# Identification of novel Rab46 effector proteins

**DOI:** 10.1101/2020.07.06.189340

**Authors:** Lucia Pedicini, Sabina D Wiktor, Katie J Simmons, Lynn McKeown

**Author notes:** Corresponding author, Address: LICAMM, Faculty of Medicine and Health, University of Leeds, Clarendon Way, Leeds. LS2 9JT. **Summary** Identification of effector proteins is key to understanding the role of Rab GTPases in cellular trafficking. We identify two Rab46 effectors in endothelial cells and provide a resource for further analysis.

## Abstract

Rab46 is a novel Ca^2+^- sensing Rab GTPase shown to have important functions in endothelial and immune cells. The presence of functional Ca^2+^- binding, coiled-coil and Rab domains suggest that Rab46 will be important for coupling rapid responses to signalling in many cell types. The molecular mechanisms underlying Rab46 function are currently unknown. Here we provide the first resource for studying Rab46 interacting proteins. Using liquid chromatography mass spectrometry (LC-MS/MS) to identify affinity purified proteins that bind to constitutively active GFP-Rab46 or inactive GFP-Rab46 expressed in endothelial cells, we have revealed 922 possible interacting proteins. Further comparative analyses between nucleotide-locked Rab46 proteins enriched this dataset to confidently identify 29 effector proteins. Importantly, through biochemical and imaging approaches we have validated two potential effector proteins; dynein and the Na^2+^/ K^+^ ATPase subunit alpha 1 (ATP1α1). Hence, our use of affinity purification and LC-MS/MS to identify Rab46 neighbouring proteins provides a valuable resource for detecting Rab46 effector proteins and analysing Rab46 functions.

## Introduction

Rab proteins are the largest member (65 in human) of the Ras superfamily of small guanosine tri-phosphatases (GTPases) (Colicelli, 2004). Rabs are master regulators of intracellular vesicle formation, transport and fusion and their importance is evident by the many human diseases caused by mutations that affect these functions (Aridor and Hannan, 2000; Stenmark, 2009). Similar to other GTPases, Rabs cycle between a GDP-bound OFF state and a GTP-bound ON state (Pfeffer, 2017). However, most Rabs lack the inherent ability to efficiently hydrolyse GTP and therefore require guanine nucleotide exchange factors (GEFs) and GTPase-activating proteins (GAPs) to regulate their nucleotide binding status (Bos et al., 2007; Novick, 2016). In this way, Rabs act as molecular switches that mediate downstream events by interacting with effector molecules when anchored, in their GTP-bound form, to their target membrane compartment.

Novel, large Rab GTPases have recently been described that, in addition to having the highly conserved C-terminal Rab domain, they also contain a coiled-coil domain and distinct N-terminal EF-hand domains that have the ability to bind Ca^2+^. To date only three Ca^2+^-sensing GTPases (Rab44; Rab45 (RASEF) and Rab46 (CRACR2A)) have been reported. These Rabs play roles in trafficking events in osteoclasts (Rab44) (Yamaguchi et al., 2018), cancer cells (Rab45) (Nakamura et al., 2011), endothelial cells (ECs) and immune cells (Rab46) (Miteva et al., 2019; Srikanth et al., 2016). Their ability to sense changes in intracellular Ca^2+^ implies that not only do these proteins have the ability to regulate intracellular trafficking events, but they could provide spatial and temporal regulation in response to signalling events (allowing a Ca^2+^ sensor to be in the right place at the right time) thereby providing a rapid link between intracellular signalling and vesicular transport.

We discovered Rab46 (CRACR2A-L) in ECs (Wilson et al., 2015) and described its important function in coupling stimuli to the appropriate release of cargo from endothelial-specific storage organelles (Weibel-Palade bodies: WPBs) (Miteva et al., 2019). In response to histamine, but not thrombin, Rab46 diverts a subpopulation of WPBs, carrying cargo superfluous to a histamine (immune) response, away from the plasma membrane to the microtubule organising centre (MTOC), inhibiting cargo release and thus preventing an all -out thrombotic response. In T-cells, Rab46 is a component of sub-synaptic vesicles and important for activation of the Ca^2+^ and the Jnk signalling pathways upon T-cell receptor stimulation (Srikanth et al., 2016). The expression profile of Rab46 suggests that it will also have importance in a wider cellular context and thereby it is crucial that we understand the molecular machinery underlying its function. Active (GTP-bound) Rab GTPases are localized to their specific membrane compartment (Pfeffer and Aivazian, 2004) where they recruit various effector proteins that mediate biogenesis, transport, tethering, and fusion of membrane-bound organelles and vesicles (Stenmark and Olkkonen, 2001). Hence identifying effectors will be key in understanding the complete function of Rab46. In ECs we have shown that the retrograde trafficking of WPBs is dependent on the GTP-bound state of Rab46 and is mediated by dynein-dependent movement along microtubules. Indeed, Wang *et al*. have suggested that Rab46 may be a Ca^2+^- dependent adaptor for dynein in T-cells (Wang et al., 2019). However, we still do not know if dynein is a direct Rab46 effector protein in ECs or what other proteins regulate Rab46 recruitment and function.

Here, we provide for the first time, a non-biased screen of Rab46 protein-protein interactors by using affinity chromatography and liquid chromatography mass spectrometry (LC-MS/MS: see Fig. 1A). In total, 922 significantly interacting proteins were identified and are provided as an appendix as a resource for further analysis. To specifically identify high confidence effector proteins, we ranked peptides pulled down with GTP-locked active Rab46 (Q604L) against proteins that interacted with a GDP-locked inactive Rab46 (N658I). This enrichment generated 29 candidate effectors with GO terms associated with membrane trafficking. Importantly, we validated the interaction of two of these proteins with endogenous Rab46 in ECs. First, we support a role for Rab46 as an adaptor for dynein (which we have already shown is necessary for Rab46-dependent trafficking (Miteva et al., 2019)), and we suggest this direct interaction potentially acts through two alanine residues in the Rab46 coiled-coil domain which are highly conserved in other dynein adaptors (Olenick and Holzbaur, 2019). Furthermore, we validate this resource tool by confirming interaction of endogenous Rab46 with a newly identified protein partner, ATP1α1.

**Figure 1:**
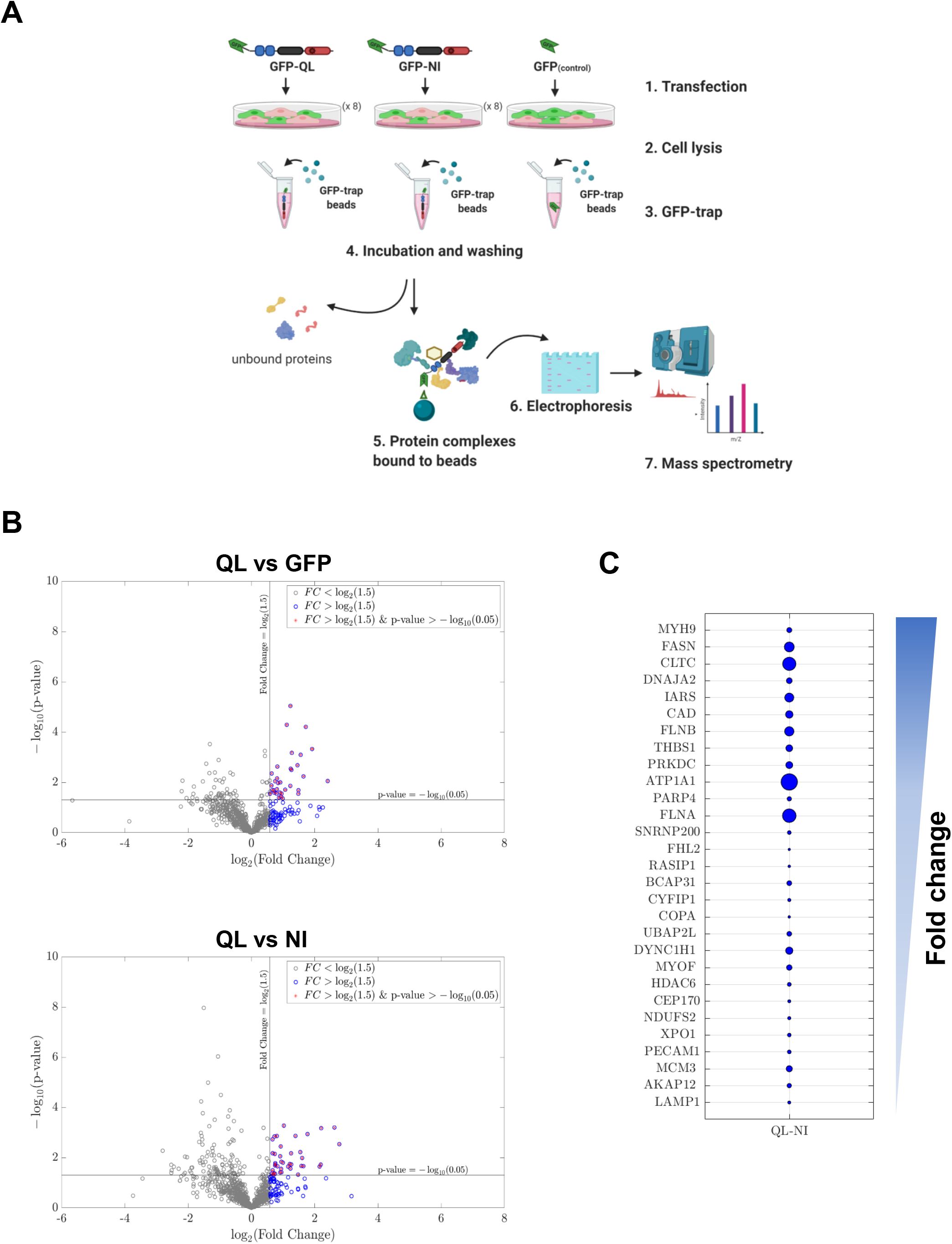
Identification of Rab46 interactome in endothelial cells. A) A schematic showing the workflow for the pull-down experiments demonstrating the major steps for purification of GFP-tagged Rab46 mutants and interacting proteins. ECs cells transfected with GFP-tagged Rab46 mutants were lysed and incubated with GFP-trap beads. After incubation the beads were sedimented and following several wash steps the protein complex bound to the beads was eluted with appropriate buffer. 10% of the beads were used for SDS-PAGE/Coomassie staining and western blot analysis. Proteins interacting with the bait protein co-purify on the affinity beads and are subsequently identified using mass spectrometry. B) Volcano plots showing the distribution of the significant proteins that co-precipitated with GFP-tagged Rab46 nucleotide binding active mutant (Q604L) versus GFP control, or versus inactive N658I mutant. Compared values of significant peptides identified from mass spectrometry analysis were plotted according to p-value (−log_10_ transformed) on the y-axis and fold change (log_2_ transformation) on the x-axis. Significance level is indicated with a horizontal straight black line (p-value < 0,05) and fold change threshold with vertical black line (fold change ≥ 1.5). Blue datapoints represent a first sorting filter to highlight proteins with a fold change greater than 1.5 and purple data points highlight proteins with also a significant p-value. C) Proteins co-precipitate with GFP-Q604L in reference to GFP-N658I, were sorted by fold change (all greater than 1.5, descending values as per arrow) and visualized using a bubble plot where the circle area is proportional to p-values (bigger circle = lower p-value).

We propose a first map of novel candidate proteins which may open up new routes in understanding Rab46 mechanism of action in ECs and potentially in a wider cellular context. Characterization of novel key proteins in the Rab46 network could allow the identification of new targets to be exploited as modulator of its physiological function.

## Results

### Identification of Rab46-interacting proteins in endothelial cells

To understand the molecular mechanism underlying Rab46 function in ECs, we sought to identify interacting protein partners. To isolate potential Rab46 effector proteins, we considered the constitutively active GTP-bound mutant (Q604L) compared to the inactive GDP-bound mutant (N658I) for proteomic analysis, as most known Rab effector proteins preferentially interact with the GTP-bound form of the Rab GTPase. We performed affinity chromatography, followed by LC-MS/MS to define proteins that bound to the Q604L and the N658I GFP-Rab46 mutants. GTPase deficiency of both the mutants has been previously validated (Srikanth et al., 2016). Fig. 1A illustrates the general workflow for the GFP-trap pull down and protein identification by mass spectrometry. Quantitative data extraction for all identified protein was performed in PeakView and peptides were statistically filtered using false discovery rate (FDR) < 1%.

Pulldown experiments and LC-MS/MS identified a total of 922 unique peptides that significantly co-purified with active Q604L, inactive N658I or GFP control. All the identified peptides are derived from proteins that are potential novel Rab46 interactors and thus, this dataset provides a valuable resource for studying Rab46 biology. However, to extricate proteins for further analysis, we enriched for likely Rab46 effectors by pairwise comparisons. This comparative analyses generated three different datasets: i) QL vs GFP, ii) NI vs GFP, iii) QL vs NI (Tables 1, 2). To rank these candidate effectors with confidence we firstly considered that true interacting proteins would bind with a higher specificity to GFP-Rab46 mutants than GFP alone, hence we extracted Q604L bound peptides that displayed a significant (p-value ≤ 0.05) 1.5-fold change in abundance from peptides that bound GFP (volcano plot Fig. 1B top). We then assumed that effector proteins would display increased binding specificity to the active Q604L form than the inactive N658I mutant, so we applied the same criteria to extract Q604L abundant bound peptides compared to N658I bound peptides (volcano plot Fig. 1B bottom). We identified 29 Q604L enriched proteins (3.5% of total proteins) with significant p-value compared to N658I peptides and 34 Q604L enriched proteins compared to GFP. Finally, in Fig. 1C (and Fig. S1) we ranked these Q604L bound proteins according to their fold change with their respective p-value represented by bubble size (bigger circle represents smaller p values). Co-purified impurities such as keratin and trypsin were removed from this list, as well as typical nonspecific binders such as heat shock proteins and elongation factors (Hodge et al., 2013).

The differentially expressed proteins in Q604L vs N658I (n = 29) were investigated for biological function using STRING database (https://string-db.org/) to establish known and predicted protein interactions (Fig. 2A). The gene ontology (GO) classification system was used to elucidate biological processes, cellular components and molecular functions of target proteins. GO terms were submitted to REVIGO (http://revigo.irb.hr/) and the results visualised using interactive plots (Fig 2B, S2) to help reveal identified patterns within the data in a comprehensive and biologically meaningful manner. Interestingly, Fig. 2B shows a central node for transport processes with different edges pointing at more specific annotation such as localization, secretion, vesicle-mediated transport and microtubule-based transport. Therefore, clear clusters emerge within the biological process for regulation of trafficking events. Moreover, analysis revealed highly represented GO cellular component terms from intracellular organelles including cytoplasmic vesicles, secretory granules and endocytic vesicles as well as cytoskeleton components (Fig. S2). GO molecular function terms have also been considered showing many binding function terms, including different subsets, to be the most abundant (Table 3). Altogether the function enrichment analysis showed a highly proportion of terms associated with membrane and vesicular trafficking processes which are of particular interest for understanding the role of Rab46 in ECs.

**Figure 2:**
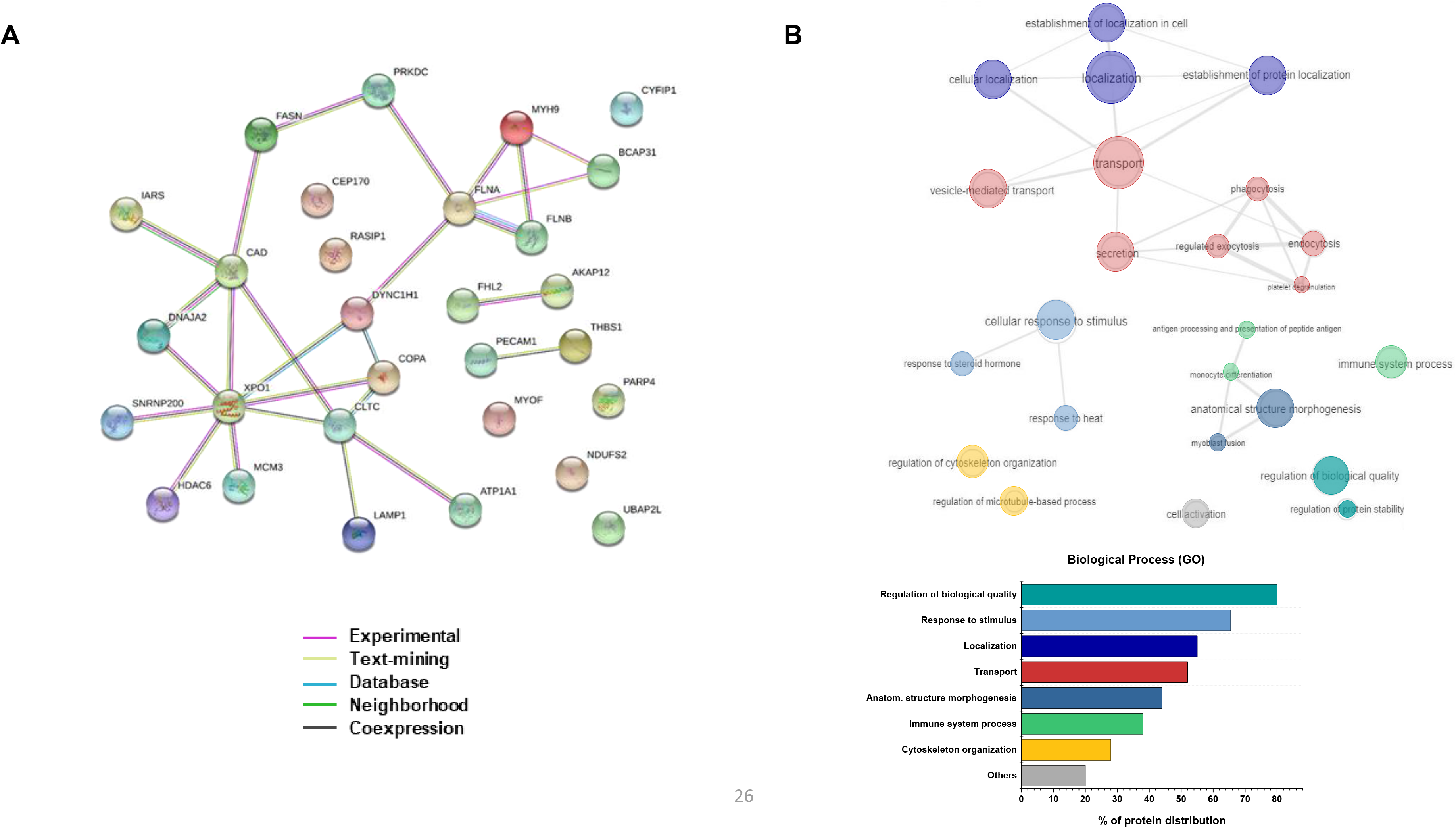
Functional profiling of potentially new Rab46 interacting partners. A) STRING PPI network connectivity of 29 proteins identified in the QL vs NI dataset. Network contains 29 nodes, 26 edges (vs 12 expected edges), 1.8 average node degree, 0.58 avg. local clustering coefficient. PPI enrichment p-value 0.000254. Different colors of the lines represent the types of evidence used in predicting the association. B) GO enrichment analysis elucidating enriched biological process (BP). GO terms identified using dataset imported from STRING and analyzed with REVIGO web server. Redundant GO terms were excluded for clarity. Results are visualized with REVIGO interactive maps sho wing enriched BP GO terms. Similar GO terms are color coded. The color matches the bar colors of the bar chart on the right showing percentage of protein distribution among the most represented GO terms identified.

### Rab46 interacts directly with the dynein complex

Among the set of high confidence Rab46 interactors identified by proteomic analysis, the dynein heavy chain (DHC) constitutes a well-known motor protein involved in cargo trafficking, therefore most likely to be a relevant physiological interactor for a small Rab GTPase. Our previous findings revealed a dynein-dependent retrograde trafficking of WPBs to the MTOC that required Rab46 GTPase activity (Miteva et al., 2019). Moreover, the identification of Rab46 as a new dynein adaptor protein in immune cells by Wang *et al*., (Wang et al., 2019) advocates DHC as a promising candidate for further investigation.

We have previously validated the interaction between Rab46 and a complex containing DHC in ECs (Miteva et al., 2019). To determine if this is a direct interaction or via adaptor proteins we considered that dynein adaptors form a complex with dynein and dynactin. The largest component of the dynactin complex is p150^Glued^ which binds to dynein and mediates direct association between dynactin and microtubules (Schroer, 1994). Co-immunoprecipitation (Co-IP) resulted in pull down of endogenous Rab46 from histamine treated ECs and demonstrated the interaction between Rab46 and the dynein-dynactin complex via both DHC and p150^Glued^ (Fig. 3A). Reverse immunoprecipitation (IP) also verified complex formation in ECs (Fig 3B-C). Probing for these proteins simultaneously (p150, Rab46 and DHC) revealed a specific band at the expected molecular weight of endogenous Rab46 (95 kDa) in both the DHC and p150 IP samples (Fig. 3B-C). The association of Rab46, DHC and p150 suggested the existence of a Rab46-dynein-dynactin motor complex, in a manner similar to other dynein adaptors. Following histamine stimulation of ECs, we also observed an increase in the Co-IP of dynein by the anti-Rab46 and p150 antibodies compared with a non-treated cells. This observation raises the question whether histamine stimulation, which we suggest switches Rab46 in the GTP-bound form, enhances dynein-dynactin complex stability.

**Figure 3:**
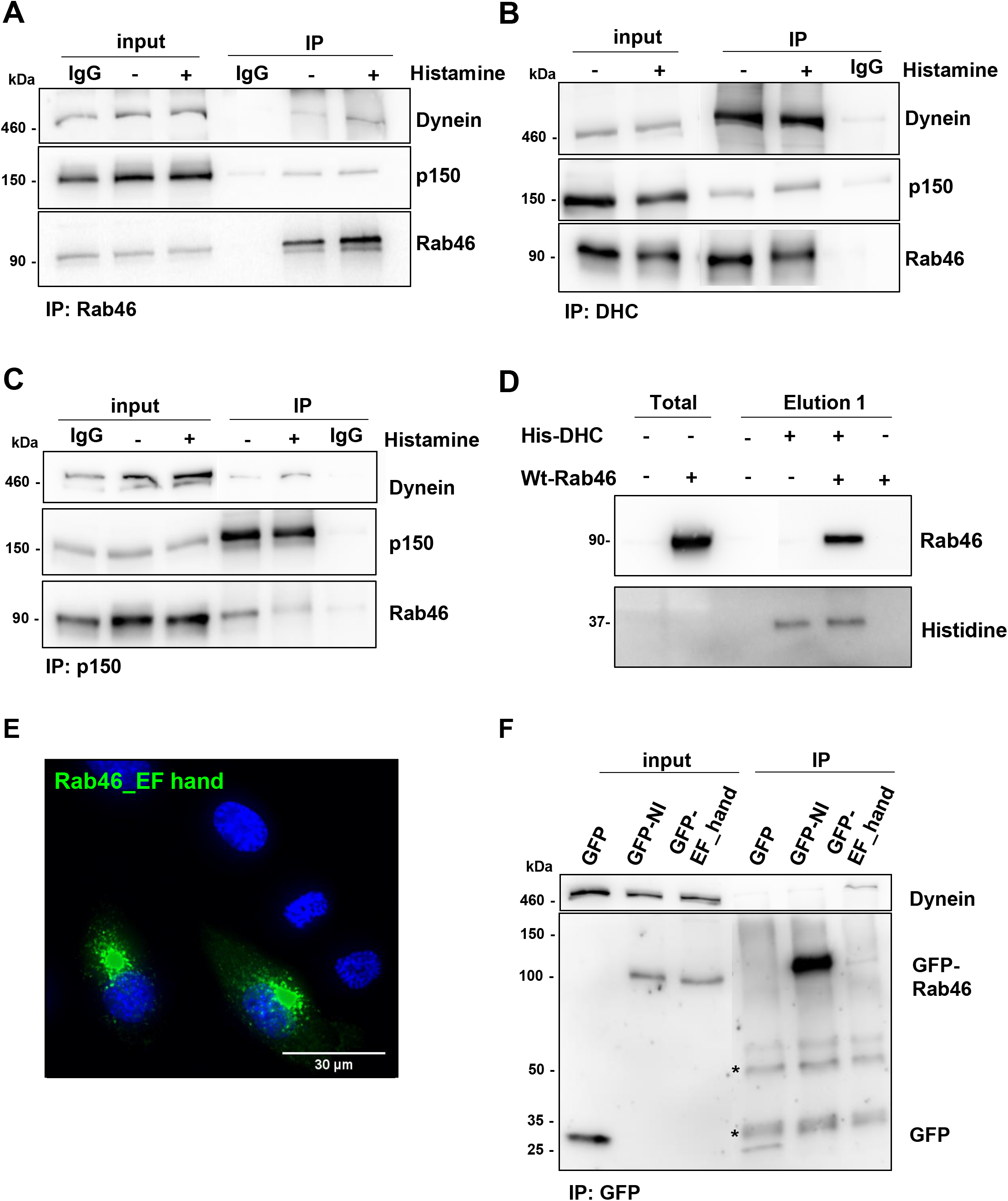
Rab46 interacts with the dynein/dynactin complex in a Ca^2+^ independent manner. A) Immunoprecipitation (IP) of endogenous Rab46 in HUVECs stimulated with 30 μM histamine or vehicle control was performed using an anti-Rab46 antibody and co-precipitation of DHC and p150, subunit of dynactin, was assessed by immunoblotting. B) Reverse Co-IP of endogenous Rab46 and p150 subunit was performed using anti-DHC antibody, co-precipitation was assessed by immunoblotting for Rab46 and p150. C) Reverse Co-IP of endogenous Rab46 and DHC was performed using anti-p150 antibody, Co-IP was assessed by immunoblotting. Input represents total lysate before the IP and samples after the IP are denoted as IP. Corresponding IgG were used as negative control. Blots are representative of 3 independent experiments. D) Pull down of human WT-Rab46 overexpressed in Cos7 cells using purified his-tag DHC (His-DHC) bound to Ni^2+^ beads. Non-transfected cells (−) and cells transfected with WT-Rab46 (+) were mixed with (+) or without (−) 10 μg of His-DHC. Magnetic sepharose Ni^2+^ beads were used to pull-down His-tag DHC complexes: the beads were incubated with the mixture of His-DHC + WT-Rab46 (or non-transfected cell as control) or WT-Rab46 only as control to show non-specific binding to the beads. Pull down of Rab46 with his-DHC was assessed by western blot using anti-Rab46 and anti-histidine antibodies. Western blots are representative of 3 independent experiments. E) Representative images of HUVECs expressing Rab46 EF-hand mutant (green). Deficiency in Ca^2+^ binding shows Rab46 EF-hand mutant clustering in the perinuclear area. DAPI (blue) represents nuclei. Scale bar = 30 μm. F) Immunoprecipitation of Rab46 binding mutants (GFP-N658I and GFP-EF hand mutant) in HUVECs was performed using an anti-GFP antibody and Co-IP of DHC was assessed by immunoblotting for DHC. Input represents total lysate before the IP and samples after the IP are denoted as IP. Empty GFP vector was used as negative control. Asterisks (*) indicates non-specific bands corresponding to the IgG heavy (50 kDa) and light chains (25 kDa).

We sought to determine whether Rab46 could directly interact with dynein. To this end, we ectopically expressed human Rab46 (hRab46) in Cos-7 (monkey) cells and used a purified recombinant portion of the tail domain of human DHC (His-DYNCH1) as a bait to identify binding partners. These cells do not express hRab46 and therefore present as a suitable system to monitor direct interactions. Western blot analysis (Fig. 3D), revealed the presence of recombinant dynein (37kDa) in the positive control and in the elution fraction after incubation with lysates containing hRab46. Probing for Rab46 revealed a single band at the expected molecular weight (95 kDa) in the pulled down fraction. No Rab46 was detected in cells not expressing hRab46 and no cross-reactivity was observed in the negative control where WT-Rab46 was incubated with the Ni^2+^ beads but in absence of His-DHC. These results suggest that the interaction between hRab46 and DHC is a direct interaction between the two proteins.

### Rab46 / dynein interaction is independent of calcium

Rab46 contains two functional Ca^2+^ binding motifs (EF-hands) in the N-terminal (Srikanth et al., 2010). Wang *et al* recently reported that Ca^2+^ binding to the EF-hands was necessary for the interaction between Rab46 and dynein in T-cells (Wang et al., 2019). Surprisingly, in ECs, chelation of intracellular Ca^2+^ does not inhibit dynein-dependent trafficking of WPBs to the MTOC (Miteva et al., 2019). Moreover, a mutant of Rab46 that is unable to bind Ca^2+^ (Rab46^EFmut^) is recruited to WPBs that rapidly translocate to the MTOC even in the absence of stimulation, further supporting Ca^2+^ independence (Fig. 3E). In order to investigate the dependence of the Rab46-dynein interaction on Ca^2+^, an anti-GFP antibody was used to pull out proteins that interacted with GFP-tagged Rab46^EFmut^ as compared to the inactive GFP-N658I Rab46 (that does not bind to DHC) (Fig. 3F). Although the level of Rab46^EFmut^ in the IP fraction is lower than the inactive GFP-N658I mutant, immunoblotting for DHC showed a distinct band, at the correct molecular weight (460 kDa), in this fraction as compared to the N658I mutant or GFP control. These results support our previous cellular imaging studies (Miteva et al., 2019) that suggests the interaction between Rab46 and dynein is Ca^2+^- independent in ECs.

Altogether these observations in ECs support the evidence given by Wang *et al*., which describe Rab46 as a new dynein adaptor in T-cells. However, complex formation between Rab46 and dynein–dynactin is insensitive to intracellular Ca^2+^ in ECs.

### Identification of specific binding sites that enable Rab46 / dynein interaction

Rab46 is an unconventional Rab that contains a pair of EF-hands, a coiled-coil domain and a Rab GTPase domain (Fig. 4A). The domain architecture resembles some well-known dynein adaptors (Fig. 4A). A common feature of dynein adaptor proteins is the presence of long coiled-coil domains containing conserved residues that are important for interaction with dynein. Particularly, the dynein adaptor Rab11FIP3 has two EF-hands followed by coiled-coil domains (Horgan and McCaffrey, 2009) that appear structurally similar to Rab46. Comparison of the amino acid sequence of Rab46 and Rab11FIP3 by global sequence alignment, demonstrated 13.6% identity and 22.9% similarity (Fig. S3). The two proteins display high similarity within the Ca^2+^ binding motif, the coiled-coil domain and a specific Rab-binding domain at the C-terminal called FIP-RBD domain. Moreover, two conserved alanine residues responsible for dynein binding in the Rab11FIP3 sequence were identified in the Rab46 sequence (A227 and A555 highlighted in red Fig. 4B). These alanines are conserved in other dynein adaptors including BICD2, BICDR1, BICDR2, Spindly and HAP1 (Gama et al., 2017; Olenick and Holzbaur, 2019). BICD family members and other adaptor proteins, such as HAP1 and TRAK1/2 have an additional alanine residue just adjacent to the aforementioned conserved alanine residue present in the coiled-coil domain (Schlager et al., 2014) which we also found next to the A227 conserved in Rab46 (Fig. S4). Identification of A227 at the N-terminal of the Rab46 coiled-coil domain and A555 within the Rab domain suggest that these residues may be important in binding to dynein.

**Figure 4:**
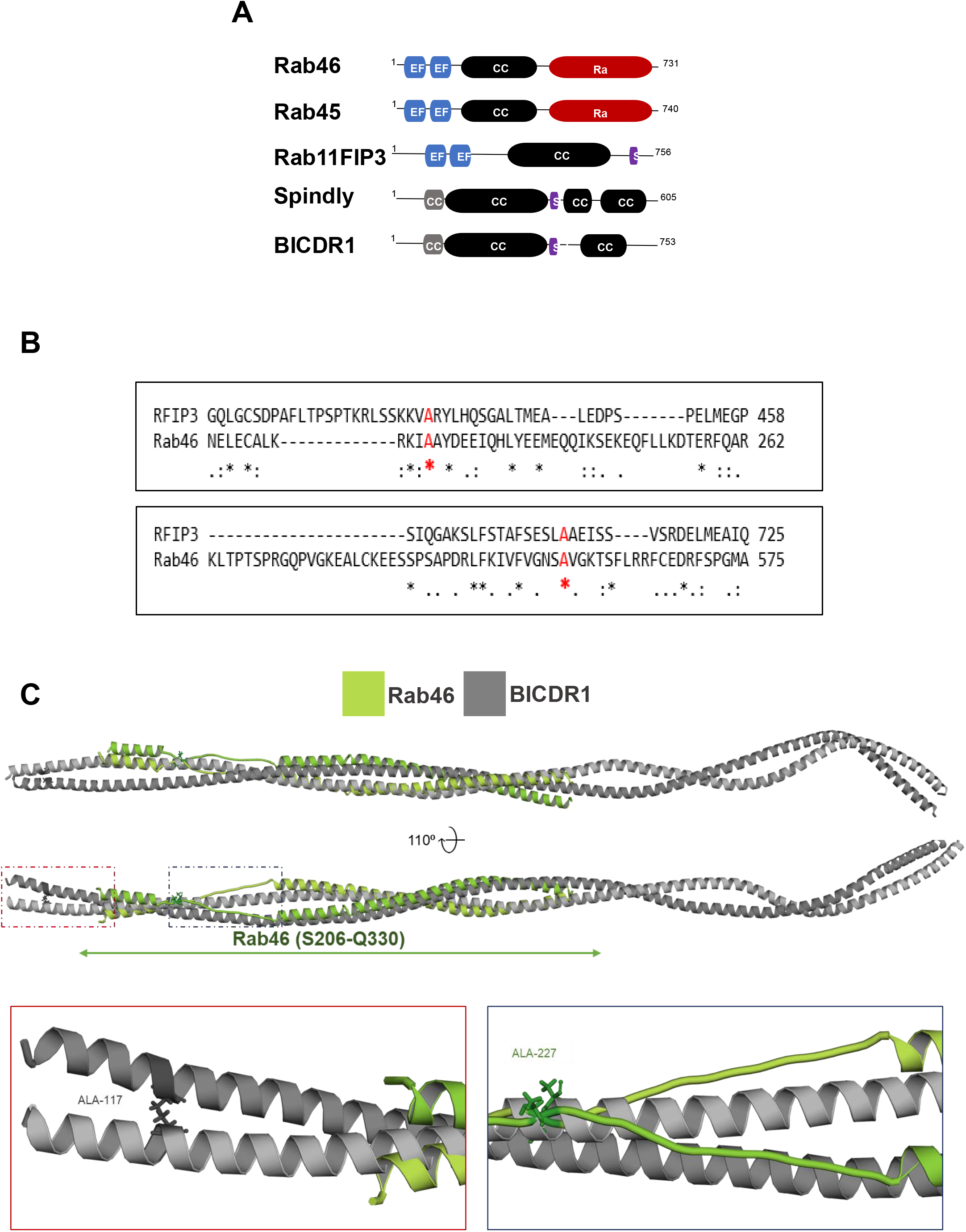
Rab46 is a novel dynein/dynactin adaptor. A) Domain organization of structural related Rab GTPases (Rab46 and Rab45) compared to Rab11FIP3, Spindly and BIC DR1 (dynein activators and adaptors). EF-hand domains (blue): calcium binding sites. Coil-coiled domain (CC: black): protein interactions sites. Rab (red): GTP/GDP binding site. CC1 (grey): coiled-coil segment shared by some dynein binding proteins. Spindly motif (S: purple): conserved features of some dynein activators. B) Sequence alignment (cropped from the full alignment showed in S2) using Clustal Ω shows a conserved region in the first coiled-coil segment of the dynein adaptor Rab11FIP3 (CC1 box) and Rab46 coiled -coil domain (top). The alignment also shows a second conserved residue in the spindly motif of RFIP3 (708 - 712) which is shared within the Rab46 sequence. The two alanine’s (A435 and A709 in RFIP3) important in the interaction with dynein and the conserved alanine’s in Rab46 sequence (A227 and A555) are marked with red asterisks. C) Homology model of Rab46 coiled-coil domain. Top: Rab46 coiled-coil structure (green) computed by the SWISS-MODEL server and superimposed on CryoEM structure of BICDR1 (grey) using PyMOL. Bottom: expanded boxes showing detailed views of the position of conserved alanine residues in BICDR1 (left) and Rab46 (right) coiled-coil domains.

Alignment of the computed Rab46 model to all structures in the PDB library identified one of the chains of BICDR1 dynein adaptor (6F1TX) as a top structural homolog of Rab46 coiled-coil (TM-score = 0.672). The alignment of Rab46 coiled-coil and BICDR1 structures demonstrated structural similarity between the proteins (reported RMSD = 4.49 Å) (Fig. 4C). Although RMSD value of less than 3 Å would typically be expected for homologous proteins (Reva et al., 1998) there are no high-resolution structures of dynein adaptor proteins currently available to be used as modelling templates. The positions of conserved alanine residues in BICDR1 and Rab46 are shown in the expanded boxes in Fig. 4C.

These preliminary results suggest Rab46 is a dynein adaptor that may interact with dynein and dynactin through conserved alanine residues found in the N-terminal domain.

### Rab46 interacts with the Na^2+^/ K^+^ ATPase subunit alpha 1

Na^2+^/ K^+^ ATPase subunit α1 (ATP1α1) was identified as another potential Rab46 effector protein in our enriched dataset. To date, the role of ATP1α1 in membrane trafficking has not been investigated. However, recently ATP1α1 has been identified as new effector for Rab27a (Booth et al., 2014). Therefore, we decided to further investigate the potential interaction between Rab46 and the ATP1α1.

Validation of the interaction between Rab46 and ATP1α1 was performed using IP and western blot analysis in lysates from cells expressing GFP-tagged constitutively active (Q604L) Rab46. A band size of 110 kDa was detected in the IP fractions and confirmed as ATP1α1. Specificity of the antibody has been validated by siRNA experiments (Fig. S5). Notably, the ATP1α1 preferentially interacts with the active GTP-bound Rab46 (Fig. 5A).

**Figure 5:**
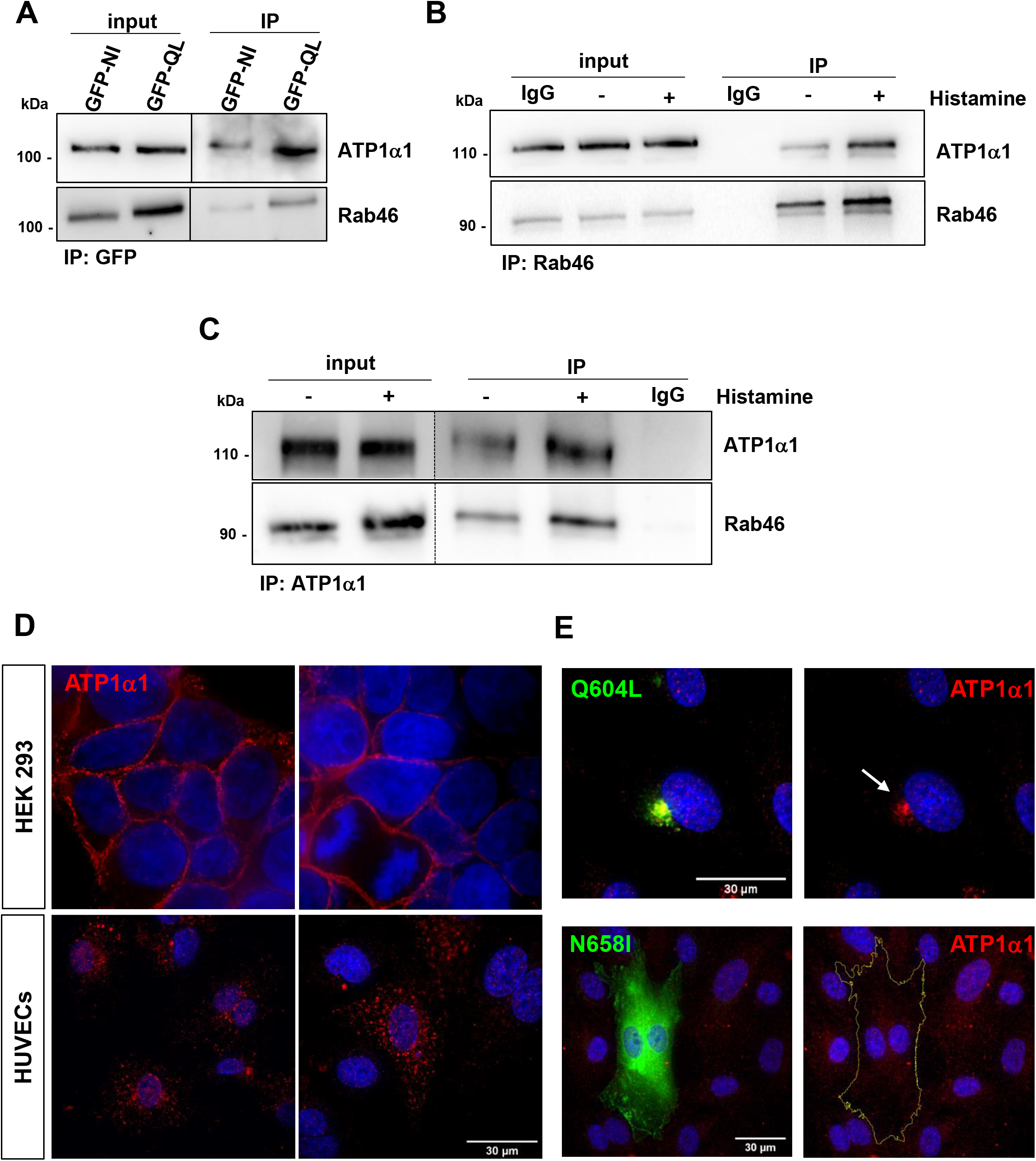
Rab46 interacts with the Na^2+^/ K^+^ ATPase subunit α1. A) IP of Rab46 nucleotide binding mutants to confirm proteomic analysis. GFP-tagged active and inactive form of Rab46 (Q604L and N658I respectively) were overexpressed in HUVECs and immunoprecipitation performed using an anti-GFP antibody. Input represents lysate before IP and IP denotes samples after IP. Western blotting using anti-GFP and anti-ATP1α1 antibodies shows that ATP1α1 Co-IPs with the active form of Rab46 (Q604L). B-C) Endogenous Rab46 interacts with ATP1α1. IP of endogenous Rab46 in HUVECs stimulated with 30 μM histamine or vehicle control was performed using an anti-Rab46 antibody and Co-IP of ATP1α1 was assessed by immunoblotting (B). Reverse Co-IP of endogenous Rab46 performed using anti-ATP1α1 antibody, Co-IP was assessed by immunoblotting for Rab46. Input represents total lysate before the IP and samples after the IP are denoted as IP (C). Anti-rabbit and anti-mouse IgG were respectively used as negative controls. Blots are representative of 3 independent experiments. D) Representative immunofluorescent images of ATP1 α1 (red) localization in HEK293 (top) and HUVECs (bottom). DAPI (blue) shows nuclei. Scale bar = 30 μm. E) Representative images of HUVECs expressing Rab46 nucleotide bindings mutants (green) : Q604L (top), N658I (bottom) and ATP1α1 (red) expression. DAPI (blue) shows nuclei. Scale bar = 30 μm.

To validate this result for the native protein, endogenous Rab46 IP was performed and ATP1α1 co-immunoprecipitation assessed by immunoblotting. To induce the active form of Rab46, ECs were treated with histamine prior to IP. Co-IP demonstrated association of endogenous Rab46 with ATP1α1 (Fig. 5B) and a stronger band is visible for ATP1α1 in the histamine-treated sample. This interaction was validated with a reverse Co-IP using an antibody against the novel target (ATP1α1) and immunoblotting for the original “bait” (Rab46). Similarly, the reverse Co-IP revealed the presence of Rab46 in both the ATP1α1 IP fractions with a stronger band corresponding to the histamine stimulated IP fraction (Fig. 5C). The specificity of this interaction is supported by the use of control IgG as negative control showing no interaction between the IgG and the target proteins.

These results suggest that ATP1α1 is a Rab46-interacting protein and that this interaction is enhanced or maybe stabilized after histamine stimulation.

### Na^2+^/ K^+^ ATPase subunit alpha 1 expression in endothelial cells

To date, the expression and the role of ATP1α1 in ECs has not been investigated. ATP1α1 was depleted in ECs by targeted siRNA and the antibody specificity validated with western blot (Fig. S5). A single band at the known molecular weight (110 kDa) was observed in the mock and control siRNA transfected cells whereas a reduced band intensity was observed in ATP1α1 siRNA transfected cells (Fig. S5). This specificity encouraged the use of this antibody for immunofluorescence imaging studies. Using high resolution imaging, we observed an intracellular vesicular-like localization of ATP1α1 in ECs (Fig. 5D), in contrast to HEK293 cells which express ATP1α1 at the plasma membrane. Moreover, the GTPase activity of Rab46 affected the intracellular localization of ATP1α1 where, ATP1α1 co-localised with the active form of Rab46 (Q604L) in the perinuclear area of ECs (Fig. 5E). Moreover, ATP1α1 displayed a cytosolic granular distribution in cells expressing the inactive N658I mutant of Rab46.

## Discussion

Rab46 is a novel Ca^2+^-sensing Rab GTPase which was first discovered in ECs by Wilson *et al* (2015). The contribution of Rab46 to the function of the endothelium (Miteva et al., 2019) and to T-cells (Srikanth et al., 2016) has recently been defined, however, the interacting proteins that regulate this function have yet to be described. In the present study we have taken an unbiased approach to identify Rab46 effector proteins by deploying label-free proteomics to generate a comprehensive list of candidate Rab46 effectors in ECs. In addition, further analyses of the candidate proteins provided a high confidence prediction of novel interactors. We have validated our findings by exploring two putative interactors; dynein and ATP1α1. Thus, here we provide a useful resource for investigating Rab46 protein-protein interactions which could serve to identify the molecular mechanisms that regulate this novel Rab GTPase.

Analysis of the 29 statistically significant enriched proteins and functional enrichment studies in our proteome dataset support the involvement of Rab46 signalling in intracellular trafficking events. Indeed, integration of functional information and topological information highlighted key proteins involved in relevant biological mechanisms. For instance, some proteins such as XPO1, DHC, FLNA, MYO9 and CYFIP1 which have already been characterized as effector proteins have been highlighted in the Rab46 network, together with proteins like COPA, CLTC, LAMP1 and ATP1α1 which appear to be enriched in transport and localization processes. Because protein function and location are often tightly linked, it was also interesting to observe that several of the enriched proteins are part of intracellular components such as organelles, vesicles and cytoskeleton parts. Therefore, the three ontology domains covered by the analysis strongly place Rab46, which is a non-conventional Rab GTPase, in the context of intracellular trafficking where direct or indirect binding with the proteins identified could lead to the regulation of different pathways in the endothelium.

In this study, we provide evidence that Rab46 directly interacts with the dynein/dynactin motor complex in ECs, corroborating the previously published evidence by Wang *et al* proposing Rab46 (CRACR2A-L) as a new dynein adaptor in T-cells (Wang et al., 2019). However, in contrast our results in ECs suggest that Rab46 interaction with dynein is Ca^2+^- independent.

Indeed, we have previously suggested that Ca^2+^ is necessary for the release of Rab46 from dynein and microtubules upon trafficking to the MTOC (Miteva et al., 2019). Although we are aware of the extensive evidence provided by Wang *et al*, this difference might reflect distinction in the physiology and/or the accessory protein expression between the two cell types. For instance, upon T-cell activation, the MTOC and Rab46 both traffic to the immunological synapse (Wang et al., 2019) whereas in ECs there is a retrograde movement of Rab46 along the microtubules towards MTOC upon stimulation (Miteva et al., 2019). In addition, whilst ECs only express Rab46 (Wilson et al., 2015), T-cells also express the short non-Rab isoform (CRACR2A-S) of the gene (*EFCAB4B*) that lacks the Rab domain (Srikanth et al., 2010). Both Rab46 and CRACR2A-S are necessary for regulating store-operated Ca^2+^ entry (SOCE) in T-cells but we found no function of Rab46 in SOCE in ECs (Wilson et al., 2015), suggesting an interplay between these two isoforms. Therefore, taken all together, we have demonstrated that, in ECs, Rab46 is a Ca^2+^- independent adaptor for dynein that mediates retrograde trafficking of vesicles (WPBs) to the MTOC upon activation.

Association of Rab proteins with motor complexes allow directional movement of various vesicular cargos along the microtubule cytoskeleton. Recent work has provided new in sights into the regulation of cytoplasmic dynein by adaptor proteins which link dynein to cargo (Fu and Holzbaur, 2014; Kardon and Vale, 2009; Reck-Peterson et al., 2018). Structural studies have also described some common feature between dynein adaptors demonstrating that the coiled-coil domains play crucial roles in dictating the interaction with dynein (Urnavicius et al., 2018, 2015). Given the Rab46 similarity with other dynein adaptors, such as Rab11FIP3, sequence alignment predicted some potential conserved binding sites within Rab46 coiled - coil domain which may be necessary for the protein-protein interaction. Domains outside of the coiled-coil region in adaptor proteins could facilitate or regulate their interactions with dynein–dynactin. Moreover, our homology modelling suggests some structural similarity between Rab46 and other dynein adaptors, indicating that manipulation of the conserved residues could further elucidate the role of Rab46 as a dynein adaptor. In addition, we have proposed (see (Miteva et al., 2019)) that, in ECs, Ca^2+^ binding to Rab46 is necessary for release from microtubules at the MTOC. It would therefore also be interesting to observe how Ca^2+^ binding to the EF-hands affects the structure of Rab46 and how this impacts on binding to the dynein complex.

Here, we have also identified and validated a second Rab46 interacting protein, the alpha 1 subunit of the Na^2+^/ K^+^ ATPase. It has been reported that ATP1α1 is expressed in a tissue- and developmental-specific manner and is primarily localised at the plasma membrane of cells such as neurons and kidney cells (Levenson, 1994; Lingrel, 1992), For the first time we reported the specific localization of ATP1α1 in resting ECs. In contrast to other cell types, we have shown that ECs present a more intracellular vesicle-like localization. Following the activation of Rab46 which leads to its perinuclear clustering, we have also shown a redistribution of ATP1α1 which clusters and co-localizes at the perinuclear area with Rab46. Translocation of membrane proteins and specific trafficking events sometimes plays an important role in their related biological function, understanding the location of ATP1α1 associated with Rab46 may be important in further defining the critical role of this novel GTPase in ECs. Interestingly, recent studies indicate that ATP1α1 also functions in activating signalling cascades independently of its ion-transport role. It is proposed that a separate pool of non-pumping Na^+^/K^+^ ATPases mediate this non-canonical function depending on its interactions with various proteins including protein and lipid kinases, membrane transporters, channels, and cellular receptors (Liang et al., 2007). Recently, for the first time, a study from Booth *et al* (Booth et al., 2014) identified a new role for the ATP1α1 as a novel Rab27a interacting protein in melanocytes where several lines of evidence strongly support an essential role for ATP1α1 in the targeting of Rab27a to melanosomes. Future studies will aim to depict whether ATP1α1 is necessary for targeting Rab46 to WPBs, in addition to regulation of WPB trafficking in ECs. Further studies are required to characterize this interaction as it should also be noted that this in vitro pull-down method detects both direct and indirect interactions.

Here, we have identified effector proteins by using a constitutively active (GTP-bound) Rab46. Differences in experimental design and tissue choice might influence our interaction network. Therefore, the choice of the biological system and the method for quantification will typically depend on the biological question of interest. For instance, we observed steric hindrance of the EF-hand by GFP-tag at the N-terminus of Rab46 (Miteva et al., 2019) therefore our approach here would not be suitable for investigating Ca^2+^-dependent proteins and events (which we suggest occur after effector engagement). Quantitative proteomics such as SILAC analysis and dynamic approaches such as proximity-dependent biotin identification (BioID) and new techniques including proximity-dependent ascorbic acid peroxidase labelling (APEX) will provide additional information allowing spatiotemporal resolution of the interactions, although these too have their limitations.

In conclusion, our study presents the first screen for Rab46 protein interaction partners in ECs with many new potential binding partners presented for further evaluation. Following up on the biological significance of novel interactors is beyond the scope of this paper. However, we point at some interesting observations that might be addressed in future studies. Finally, we lay out a rough map of Rab46 effectors that can contribute to regulation of trafficking events, providing a rich resource for the community that has so far been lacking.

## Material and Methods

### Cell culture

Pooled Human Umbilical Vein Endothelial Cells (HUVECs) (Lonza Inc, USA or Promocell, UK), grown in endothelial basal cell medium 2 supplemented with EGM-2 Singlequot supplements (Lonza). HUVECs maintained at 37°C in a humidified atmosphere of 5% CO_2_ and used between passages 1-5. Human Embryonic Kidney 293 cells (HEK293T – Invitrogen, UK) were grown in Dulbecco’s modified Eagle medium (DMEM - Invitrogen, Paisley, UK) containing D-glucose, L-glutamine, and pyruvate (Life Technologies, UK) supplemented with 10% Fetal Bovine Serum (FBS - Sigma-Aldrich), 100 U/m penicillin, and 100 mg/ml streptomycin (Sigma-Aldrich). Cos-7 cells were obtained from the American Type Culture Collection (ATCC) and maintained in DMEM supplemented with 10% FBS and 100 U/ml penicillin + 100 mg/ml streptomycin. Cells were maintained at 37°C and 5% CO_2_ in a humidified incubator and used between passage 2 and 20.

### cDNA constructs and Rab46 mutagenesis

The eGFP-C1 plasmid (kanamycin resistant; Clontech; 4731 bp) was used to generate N-terminal GFP-tagged Rab46 mutants. Various mutants of Rab46 were generated by PCR amplification (pHusion DNA polymerase) and site-directed mutagenesis using primers as described in the supplementary table in (Miteva et al., 2019)

### cDNA transfection

HUVECs were plated either into Ibidi μ-slide 8-well (7*10^4^ cells/ml) or into a 10 cm Petri dish (10*10^5^ cells/ml) and after 24 hours they were transfected using Lipofectamine ™ 2000 (Thermo Fisher Scientific). A 3:1 ratio between Lipofectamine and cDNA (100 ng and 6 μg respectively) was used. 1 hour after transfection the medium was removed and fresh cell culture medium added. Experiments were performed 24 hours post-transfection. COS-7 cells were plated into a 10 cm Petri dish (2*10^6^ cells/petri dish) and transfected after 24 hours as described above.

### siRNA transfection

Control siRNA (D-001810-01-05), used as a negative control, and On-TARGET plus SMART pools for ATP1α1 (L-006111-00-0005) were purchased from Dharmacon (Thermo Scientific). HUVECs at 80–90% confluence were used for transfection, which was performed using a 1:3 ratio of 100 nmol/L siRNAs with Lipofectamine 2000 reagent (Invitrogen) diluted in OptiMEM (Gibco) as per the manufacturers’ instructions. Cells were incubated with the transfection solution for 5 hours before medium change. Knockdown was assessed via western blotting. Experiments were performed at 72 hours after transfection.

### Western blotting

Cells were harvested with NP-40 lysis buffer (ThermoFisher Scientific, UK) with protease and phosphatase inhibitors cocktails (Sigma). Samples were loaded on 7.5% or 4-20% gels and resolved by electrophoresis. Proteins were transferred to PVDF membranes using Mini Trans - Blot^®^ Cell (BioRad). Membranes incubated for 1 hour in blocking solution consisting of 5% w/v milk diluted in TBS-T 145 mM NaCl, 20 mM Tris-base, pH 7.4, 0.5% Tween-20 and labelled with primary antibody overnight at 4°C for DHC (Proteintech – 1:1000), Rab46 (CRACR2A: Proteintech 1:800), p150^Glued^ (BD Biosciences – 1:1000), ATP1α1 (Santa Cruz – 1:500), GFP (Thermofisher – 1:1000) and histidine (BioRad – 1:1000). Immunoblots visualised using HRP-conjugated donkey anti-mouse, anti-rabbit secondary antibodies (Jackson ImmunoResearch – 1:10000) and SuperSignal Femto (Pierce).

### Immunoprecipitation

HUVECs plated in 10 cm Petri dishes were starved in serum-free M199 plus 10 mM HEPES medium (Gibco) for 1 hour before treatments. Histamine (Sigma) was used at 30 μM. After treatments cells were quickly washed with PBS and before harvesting they were cross-linked with 1% PFA and then harvested with 250 μl lysis buffer (NP-40). The lysate was quantified and 0.5 mg of total lysate was incubated with 1 μg of antibody or control IgG for 4 hours at 4°C rotating. This mixture was then added to 30 μl of pre-equilibrated Protein G sepharose beads (GE Healthcare Life Science) and incubated ON under continuous agitation. Beads were washed thoroughly before elution of the bound proteins in 20 μl 4X sample buffer solution (200 mM Tris pH 6.8, 8% SDS, 40% glycerol, 8% mercaptoethanol, 0.1% bromophenol blue) and boiled at 95°C for 5 mins. The elution fraction was loaded onto SDS-PAGE gel and the amount of Rab46, dynein heavy chain, dynactin and ATP1α1 detected by western blot.

### Pull down assay

#### GFP trap

HUVECs plated in 10 cm Petri-dishes were transfected with the appropriate GFP plasmids for 24 hours. Cells were washed with cold PBS and lysed with 250 μl NP-40 lysis buffer. Lysates were left on ice for 20 mins and then centrifuged at 12000g for 10 mins at 4°C. The supernatant was then collected and the protein content was quantified using a Bio-Rad assay. 25 μl GFP-Trap bead 50% slurry (Chromotek, Planegg-Martinsried, Germany) was used and all wash steps were performed with washing buffer containing 10 mM Tris/Cl pH 7.5, 150 mM NaCl and 0.5 mM EDTA. GFP-Trap beads were washed 3x with dilution buffer prior to addition to cell lysate. Beads were incubated with cell lysate at 4°C for 2 hours following another wash step (x3). To elute the proteins off the beads 40 μl sample buffer (200 mM Tris pH 6.8, 8% SDS, 40% glycerol, 8% mercaptoethanol, 0.1% bromophenol blue) was added and samples were boiled at 95°C for 5 mins. Western blotting was used for analysis.

#### His-tag

Cos-7 cells were plated in a 10 cm Petri dish and transfected with WT-Rab46 cDNA. 24 hours after transfection cells were washed with ice cold PBS and lysed with 400 μl EDTA-free lysis buffer (BOSTER, Pleasanton, CA, USA) plus protease/phosphates inhibitor cocktail EDTA - free 100x (ThermoFisher). Cell lysate was left on ice for 20 mins and then centrifuged at 4°C 12000 g for 10 mins. Supernatant was collected and protein quantified. 20 μg of total lysate was used to incubate with pre-equilibrated beads. 100 μl of His Mag Sepharose^®^ Ni beads (GE Healthcare) were equilibrated with equilibration buffer containing 20 mM sodium phosphate, 500 mM NaCl, and 20 mM imidazole. Immediately after equilibration, total lysate was added and incubated for 1 hour at 4°C, rotating. The pre-cleared lysate was collected and the beads discarded. At this point, 10 μg of recombinant his-tagged dynein heavy chain (DYNC1H1-CloudClone) were mixed with the cleared lysate and incubated for 1 hour. After incubation, the mixture containing the complex (WT-Rab46 + His_6_DYNC1H1), was added to 100 μl of pre-equilibrated His Mag Sepharose^®^ Ni beads and incubated for 1 hour at 4°C, rotating. At this point the supernatant was discarded and a linear gradient of imidazole (up to 90 mM) was applied to the beads to reduce unspecific binding. Following these washing steps, elution was performed with elution buffer containing 20 mM sodium phosphate, 500 mM NaCl, and 500 mM imidazole. The eluted samples were collected and analysed by western blot.

### Sample preparation for mass spectrometry

Pulldown eluates were processed using the FASP procedure (Wiśniewski et al., 2009). Samples were loaded on Vivacon 500, 30k MWCO HY filter vials (Sartorius Stedim Biotech, VN01H22) and centrifuged to concentrate the proteins on the filter. The samples were washed using FASP1 by centrifugation at 7,000g. The filters were then washed by centrifugation using FASP 2 (100 mM Tris/HCl pH 8.5) ready for trypsin digestion. The samples were then reduced using 50 mM fresh IAA in FASP 2 in the dark for 30 mins. Excess buffer was removed by centrifugation and buffer exchanged into FASP 3 (100 mM TEAB - triethyl ammonium bicarbonate). Trypsin was dissolved in FASP 3 to give a 1:200 enzyme to protein ratio and added in a volume of at least 125 μL for overnight digestion (16 hrs at 30°C). This was repeated with fresh trypsin for a 5 hr incubation. Filters were spun down using 0.5 M NaCl and 150 μl 10% TFA added to reduce the pH. A standard desalting procedure was used.(Thingholm et al., 2006). After desalting, the samples were dried in a speed vac.

### LC-MS/MS and peptide identification

Desalted tryptic peptides were dissolved in 5% acetonitrile (ACN), 0.05% TFA and separated on liquid chromatograph Eksigent Ekspert nano LC 400 (SCIEX, Dublin, CA, USA). Liquid chromatography was online connected to Triple-TOF 5600+ mass spectrometer (SCIEX, Toronto, Canada). Samples were pre-concentrated on a cartridge trap column (300 μm i.d. × 5 mm) packed with C18 PepMap100 sorbent with 5 μm particle size (Thermo Scientific, MA, USA) using a mobile phase composed from 0.05% trifluoroacetic acid (TFA) in 2% acetonitrile (ACN). Pre-concentrated peptides were separated on a capillary analytical column (75 μm i.d. × 500 mm) packed with C18 PepMap100 sorbent, 2 μm particle size (Thermo Fisher Scientific, MA, USA). Mobile phase A composed of 0.1% (v/v) formic acid (FA) in water while mobile phase B composed of 0.1% (v/v) FA in ACN. Analytical gradient started from 2% B, the proportion of mobile phase B increased linearly up to 40% B in 70 min, flow was 300 nl/min. The analytes were ionized in nano-electrospray ion source, where temperature and flow of drying gas was set to 150°C and 12 psi. Voltage at the capillary emitter was 2.65 kV.

SWATH data acquisition was done in high sensitivity mode and precursor range was set from 400 Da up to 1200 Da. It was divided to 67 precursor SWATH windows with 12 Da width and 1 Da overlap. Cycle time was 3.5 seconds.

Pooled spectral library sample was measured in information dependent mode (IDA). Precursor range was set from 400 Da up to 1250 Da in MS mode and from 200 Da up to 1600 Da in MS/MS mode. Cycle time was set to 2.3 seconds and during each cycle top 20 the most intensive precursor ions were fragmented. Precursor exclusion time was set to 12 seconds. IDA data were searched against human database in ProteinPilot 4.5 (AB-SCIEX, Canada).

Quantitative data extraction for all identified protein was performed in PeakView 1.2.0.3. The quantitative data were extracted for proteins using FDR < 1%. Quantitative data were extracted using a method with 8 minute extraction window.

### Ranking of Rab46 candidate effectors

Extracted data were analysed in MarkerView 1.2.1.1. Pairwise changes in protein levels of: i) QL vs GFP, ii) NI vs GFP, or iii) QL vs NI (raw data Tables 1, 2) across samples were determined using t-test. These comparisons (as described below) use fold change values which are calculated by comparing the peak areas and p-values for significance between the technical and biological triplicates. QL vs NI comparisons were used to identify potential effector proteins and ranked according to fold change (1.5 fold change cut-off) of and p-values < 0.05. Volcano plots used to display statistical significance (p-value) versus magnitude of change (fold change) of Rab46 (Q604L) proteomic data were generated with MATLAB. Enriched peptides which met the set criteria were ranked by fold change and visualized with bubble blot to also show the significance of each protein. Bubble size represents p -values (−log_10_ p-values), larger bubbles equal more significant p-values. Interaction networks and Gene Ontology (GO) enrichment analysis of identified proteins was performed with STRING (http://string-db.org/) using default setting. The enriched GO terms were visualized using the interactive plot with REVIGO (http://revigo.irb.hr/) which was used to remove redundant GO terms and produce graphs highlighting the similarity between the terms. Highly similar GO terms are linked by edges in the plot, where the line width indicates the degree of similarity. The initial placement of the nodes is determined by a ‘force-directed’ layout algorithm that aims to keep the more similar nodes closer together (Supek et al., 2011). OriginPro 2017 software was used for bar chart presentation.

### Immunocytochemistry

Cells seeded 8×10^4^ cells/ml into Ibidi μ-slide 8 well were grown for 48 hours. Cells were fixed with 4% PFA for 10 mins, washed with PBS and permeabilised with 0.1% Triton-X solution. Primary antibodies against Rab46 (Proteintech, 1:100) and ATP1a (SantaCruz, 1:100) were added to the cells for 1 hour at room temperature. Next, a fluorescently labelled appropriate species secondary antibodies was used for 30 mins (anti-mouse Alexa 594, anti-rabbit Alexa 488 Jackson ImmunoResearch, 1:300). Cells briefly incubated in Hoechst before being mounted with Ibidi mounting medium.

### DeltaVision wide-field deconvolution microscopy

Cells visualized on an Olympus IX-70 inverted microscope using 40x/1.35 oil objectives supported by a DeltaVision deconvolution system (Applied Precision LLC) with SoftWorx image acquisition and analysis software.10 focal planes at 0.2 μm per z-stack were taken using a Roper CoolSNAP HQ CCD camera. Iterative deconvolution (5x) was performed on z stacks using the proprietary algorithm. The filter sets used were DAPI, FITC, TRITC. All imaging performed at room temperature. All the images were processed and analysed with Fiji ImageJ.

### Homology modelling

The coiled-coil domain of Rab46 (L201-R373) was modelled as a monomer using I-TASSER server (Yang et al., 2015) for protein structure and function prediction (https://zhanglab.ccmb.med.umich.edu/I-TASSER/). Alignment of the computed model to all structures in the PDB library was performed using TM-align tool (https://zhanglab.ccmb.med.umich.edu/TM-align/). A homology model of Rab46 coiled-coil domain dimer was computed using SWISS-MODEL server (Waterhouse et al., 2018) (https://swissmodel.expasy.org/). Since sequence identity between coiled-coil domains of dynein adaptors is typically low, the CryoEM structure of mouse BICDR1 (Urnavicius et al., 2018) (PDB ID 6F1TX and 6F1Tx) was used for template-based protein structure modelling in order to obtain a parallel homodimeric model of Rab46 coiled-coil. The structures of BICDR1 (105-392) and Rab46 (206-330) coiled-coils were pre-processed and energy-minimised (Roos et al., 2019) using Protein Preparation Wizard (Maestro software, Release 2020-1, Glide, Schrödinger, LLC, New York, NY, 2020). The steric clashes generated during modelling were removed using Chiron web server(Ramachandran et al., 2011). Superimposition of the protein structures and RMSD calculations were performed using PyMOL (The PyMOL Molecular Graphics System, Version 2.0 Schrödinger, LLC.).

## Abbreviations

LC-MS/MS: liquid chromatography mass spectrometry
GEFs: liquid chromatography mass spectrometr
GAPs: GTPase activating proteins
WPBs: Weibel-Palade bodies
ECs: endothelial cells
MTOC: microtubule organising centre
FDR: false discovery rate
DHC: dynein heavy chain
ATP1α1: Na^2+^/ K^+^ ATPase subunit α1

## ACKNOWLEDGEMENTS

We thank Bio-imaging Facility staff (Leeds) for their advice and support. Thank you to Gemini Biosciences for the LC-MS/MS analysis. Special thanks to Dr Niamh Forde, Mr. Ashley Money and Mrs. Katarina Miteva for reading the manuscript.

## AUTHOR CONTRIBUTIONS

L.M. conceived the study, designed experiments, analyzed the results. L.P designed experiments, performed all pull down experiments, western blotting experiments, immunofluorescent imaging and analyzed results. S.W and K.S performed homology modelling experiments. L.P. and L.M. wrote the paper. All authors reviewed the results and approved the final version of the manuscript.

## REFERENCES

Aridor, M., Hannan, L.A., 2000. Traffic Jam: A Compendium of Human Diseases that Affect Intracellular Transport Processes. Traffic 1, 836–851. https://doi.org/10.1034/j.1600-0854.2000.011104.x

Booth, A.E.G., Tarafder, A.K., Hume, A.N., Recchi, C., Seabra, M.C., 2014. A Role for Na+,K+-ATPase α1 in Regulating Rab27a Localisation on Melanosomes. PLoS One 9. https://doi.org/10.1371/journal.pone.0102851

Bos, J.L., Rehmann, H., Wittinghofer, A., 2007. GEFs and GAPs: Critical Elements in the Control of Small G Proteins. Cell 129, 865–877. https://doi.org/10.1016/j.cell.2007.05.018

Colicelli, J., 2004. Human RAS superfamily proteins and related GTPases. Sci. STKE 2004, RE13. https://doi.org/10.1126/stke.2502004re13

Fu, M., Holzbaur, E.L.F., 2014. Integrated regulation of motor-driven organelle transport by scaffolding proteins. Trends Cell Biol 24, 564–574. https://doi.org/10.1016/j.tcb.2014.05.002

Gama, J.B., Pereira, C., Simões, P.A., Celestino, R., Reis, R.M., Barbosa, D.J., Pires, H.R., Carvalho, C., Amorim, J., Carvalho, A.X., Cheerambathur, D.K., Gassmann, R., 2017. Molecular mechanism of dynein recruitment to kinetochores by the Rod–Zw10–Zwilch complex and Spindly. J Cell Biol 216, 943–960. https://doi.org/10.1083/jcb.201610108

Hodge, K., Have, S.T., Hutton, L., Lamond, A.I., 2013. Cleaning up the masses: Exclusion lists to reduce contamination with HPLC-MS/MS. J Proteomics 88, 92–103. https://doi.org/10.1016/j.jprot.2013.02.023

Horgan, C.P., McCaffrey, M.W., 2009. The dynamic Rab11-FIPs. Biochem Soc Trans 37, 1032–1036. https://doi.org/10.1042/BST0371032

Kardon, J.R., Vale, R.D., 2009. Regulators of the cytoplasmic dynein motor. Nat Rev Mol Cell Biol 10, 854–865. https://doi.org/10.1038/nrm2804

Levenson, R., 1994. Isoforms of the Na,K-ATPase: Family members in search of function, in: Reviews of Physiology, Biochemistry and Pharmacology, Volume 123: Volume: 123, Reviews of Physiology, Biochemistry and Pharmacology. Springer, Berlin, Heidelberg, pp. 1–45. https://doi.org/10.1007/BFb0030902

Liang, M., Tian, J., Liu, L., Pierre, S., Liu, J., Shapiro, J., Xie, Z. -J., 2007. Identification of a pool of non-pumping Na/K-ATPase. J. Biol. Chem. 282, 10585–10593. https://doi.org/10.1074/jbc.M609181200

Lingrel, J.B., 1992. Na,K-ATPase: isoform structure, function, and expression. J. Bioenerg. Biomembr. 24, 263–270. https://doi.org/10.1007/BF00768847

Miteva, K.T., Pedicini, L., Wilson, L.A., Jayasinghe, I., Slip, R.G., Marszalek, K., Gaunt, H. J., Bartoli, F., Deivasigamani, S., Sobradillo, D., Beech, D.J., McKeown, L., 2019. Rab46 integrates Ca2+ and histamine signaling to regulate selective cargo release from Weibel-Palade bodies. J. Cell Biol. 218, 2232–2246. https://doi.org/10.1083/jcb.201810118

Nakamura, S., Takemura, T., Tan, L., Nagata, Y., Yokota, D., Hirano, I., Shigeno, K., Shibata, K., Fujie, M., Fujisawa, S., Ohnishi, K., 2011. Small GTPase RAB45-mediated p38 activation in apoptosis of chronic myeloid leukemia progenitor cells. Carcinogenesis 32, 1758–1772. https://doi.org/10.1093/carcin/bgr205

Novick, P., 2016. Regulation of membrane traffic by Rab GEF and GAP cascades. Small GTPases 7, 252–256. https://doi.org/10.1080/21541248.2016.1213781

Olenick, M.A., Holzbaur, E.L.F., 2019. Dynein activators and adaptors at a glance. J Cell Sci 132, jcs227132. https://doi.org/10.1242/jcs.227132

Pfeffer, S., Aivazian, D., 2004. Targeting Rab GTPases to distinct membrane compartments. Nat. Rev. Mol. Cell Biol. 5, 886–896. https://doi.org/10.1038/nrm1500

Pfeffer, S.R., 2017. Rab GTPases: master regulators that establish the secretory and endocytic pathways. Mol Biol Cell 28, 712–715. https://doi.org/10.1091/mbc.E16-10-0737

Ramachandran, S., Kota, P., Ding, F., Dokholyan, N.V., 2011. Automated Minimiza tion of Steric Clashes in Protein Structures. Proteins 79, 261–270. https://doi.org/10.1002/prot.22879

Reck-Peterson, S.L., Redwine, W.B., Vale, R.D., Carter, A.P., 2018. The Cytoplasmic Dynein Transport Machinery and its Many Cargoes. Nat Rev Mol Cell Biol 19, 382–398. https://doi.org/10.1038/s41580-018-0004-3

Reva, B.A., Finkelstein, A.V., Skolnick, J., 1998. What is the probability of a chance prediction of a protein structure with an rmsd of 6 A? Fold Des 3, 141–147. https://doi.org/10.1016/s1359-0278(98)00019-4

Roos, K., Wu, C., Damm, W., Reboul, M., Stevenson, J.M., Lu, C., Dahlgren, M.K., Mondal, S., Chen, W., Wang, L., Abel, R., Friesner, R.A., Harder, E.D., 2019. OPLS3e: Extending Force Field Coverage for Drug-Like Small Molecules. J. Chem. Theory Comput. 15, 1863–1874. https://doi.org/10.1021/acs.jctc.8b01026

Schlager, M.A., Serra-Marques, A., Grigoriev, I., Gumy, L.F., Esteves da Silva, M., Wulf, P.S., Akhmanova, A., Hoogenraad, C.C., 2014. Bicaudal D Family Adaptor Proteins Control the Velocity of Dynein-Based Movements. Cell Reports 8, 1248–1256. https://doi.org/10.1016/j.celrep.2014.07.052

Schroer, T.A., 1994. Structure, function and regulation of cytoplasmic dynein. Current Opinion in Cell Biology 6, 69–73. https://doi.org/10.1016/0955-0674(94)90118-X

Srikanth, S., Jung, H.-J., Kim, K.-D., Souda, P., Whitelegge, J., Gwack, Y., 2010. A novel EF-hand protein, CRACR2A, is a cytosolic Ca2+ sensor that stabilizes CRAC channels in T cells. Nat. Cell Biol. 12, 436–446. https://doi.org/10.1038/ncb2045

Srikanth, S., Kim, K.-D., Gao, Y., Woo, J.S., Ghosh, S., Calmettes, G., Paz, A., Abramson, J., Jiang, M., Gwack, Y., 2016. A large Rab GTPase encoded by CRACR2A is a component of subsynaptic vesicles that transmit T cell activation signals. Sci Signal 9, ra31. https://doi.org/10.1126/scisignal.aac9171

Stenmark, H., 2009. Rab GTPases as coordinators of vesicle traffic. Nat. Rev. Mol. Cell Biol. 10, 513–525. https://doi.org/10.1038/nrm2728

Stenmark, H., Olkkonen, V.M., 2001. The Rab GTPase family. Genome Biology 2, reviews3007.1. https://doi.org/10.1186/gb-2001-2-5-reviews3007

Supek, F., Bošnjak, M., Škunca, N., Šmuc, T., 2011. REVIGO Summarizes and Visualizes Long Lists of Gene Ontology Terms. PLOS ONE 6, e21800. https://doi.org/10.1371/journal.pone.0021800

Thingholm, T.E., Jørgensen, T.J.D., Jensen, O.N., Larsen, M.R., 2006. Highly selective enrichment of phosphorylated peptides using titanium dioxide. Nat Protoc 1, 1929–1935. https://doi.org/10.1038/nprot.2006.185

Urnavicius, L., Lau, C.K., Elshenawy, M.M., Morales-Rios, E., Motz, C., Yildiz, A., Carter, A.P., 2018. Cryo-EM shows how dynactin recruits two dyneins for faster movement. Nature 554, 202–206. https://doi.org/10.1038/nature25462

Urnavicius, L., Zhang, K., Diamant, A.G., Motz, C., Schlager, M.A., Yu, M., Patel, N.A., Robinson, C.V., Carter, A.P., 2015. The structure of the dynactin complex and its interaction with dynein. Science 347, 1441–1446. https://doi.org/10.1126/science.aaa4080

Wang, Y., Huynh, W., Skokan, T.D., Lu, W., Weiss, A., Vale, R.D., 2019. CRACR2a is a calcium-activated dynein adaptor protein that regulates endocytic traffic. J Cell Biol 218, 1619–1633. https://doi.org/10.1083/jcb.201806097

Waterhouse, A., Bertoni, M., Bienert, S., Studer, G., Tauriello, G., Gumienny, R., Heer, F.T., de Beer, T.A.P., Rempfer, C., Bordoli, L., Lepore, R., Schwede, T., 2018. SWISS - MODEL: homology modelling of protein structures and complexes. Nucleic Acids Res. 46, W296–W303. https://doi.org/10.1093/nar/gky427

Wilson, L.A., McKeown, L., Tumova, S., Li, J., Beech, D.J., 2015. Expression of a long variant of CRACR2A that belongs to the Rab GTPase protein family in endothelial cells. Biochem. Biophys. Res. Commun. 456, 398–402. https://doi.org/10.1016/j.bbrc.2014.11.095

Wiśniewski, J.R., Zougman, A., Nagaraj, N., Mann, M., 2009. Universal sample preparation method for proteome analysis. Nature Methods 6, 359–362. https://doi.org/10.1038/nmeth.1322

Yamaguchi, Y., Sakai, E., Okamoto, K., Kajiya, H., Okabe, K., Naito, M., Kadowaki, T., Tsukuba, T., 2018. Rab44, a novel large Rab GTPase, negatively regulates osteoclast differentiation by modulating intracellular calcium levels followed by NFATc1 activation. Cell. Mol. Life Sci. 75, 33–48. https://doi.org/10.1007/s00018-017-2607-9

Yang, J., Yan, R., Roy, A., Xu, D., Poisson, J., Zhang, Y., 2015. The I-TASSER Suite: protein structure and function prediction. Nature Methods 12, 7–8. https://doi.org/10.1038/nmeth.3213

