## Supplementary figures for "Identification of novel Rab46 effector proteins"

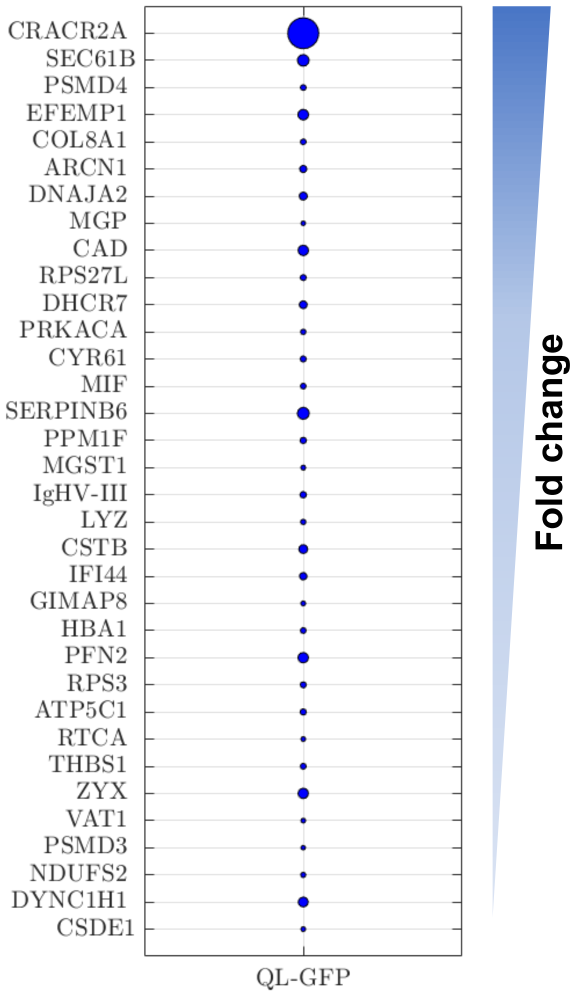


**Fig. S1: Rab46 enriched proteins compared to GFP control.** Proteins co-precipitate with GFP-Q604L identified by mass spectrometry analysis in reference to GFP control with fold change ≥ 1.5 and significant p-value. Enriched protein are ranked based on changes of fold change and the p-value visualised with a bubble plot where the circle area is proportional to p-values. Bigger circles show smaller (more significant) p-values.

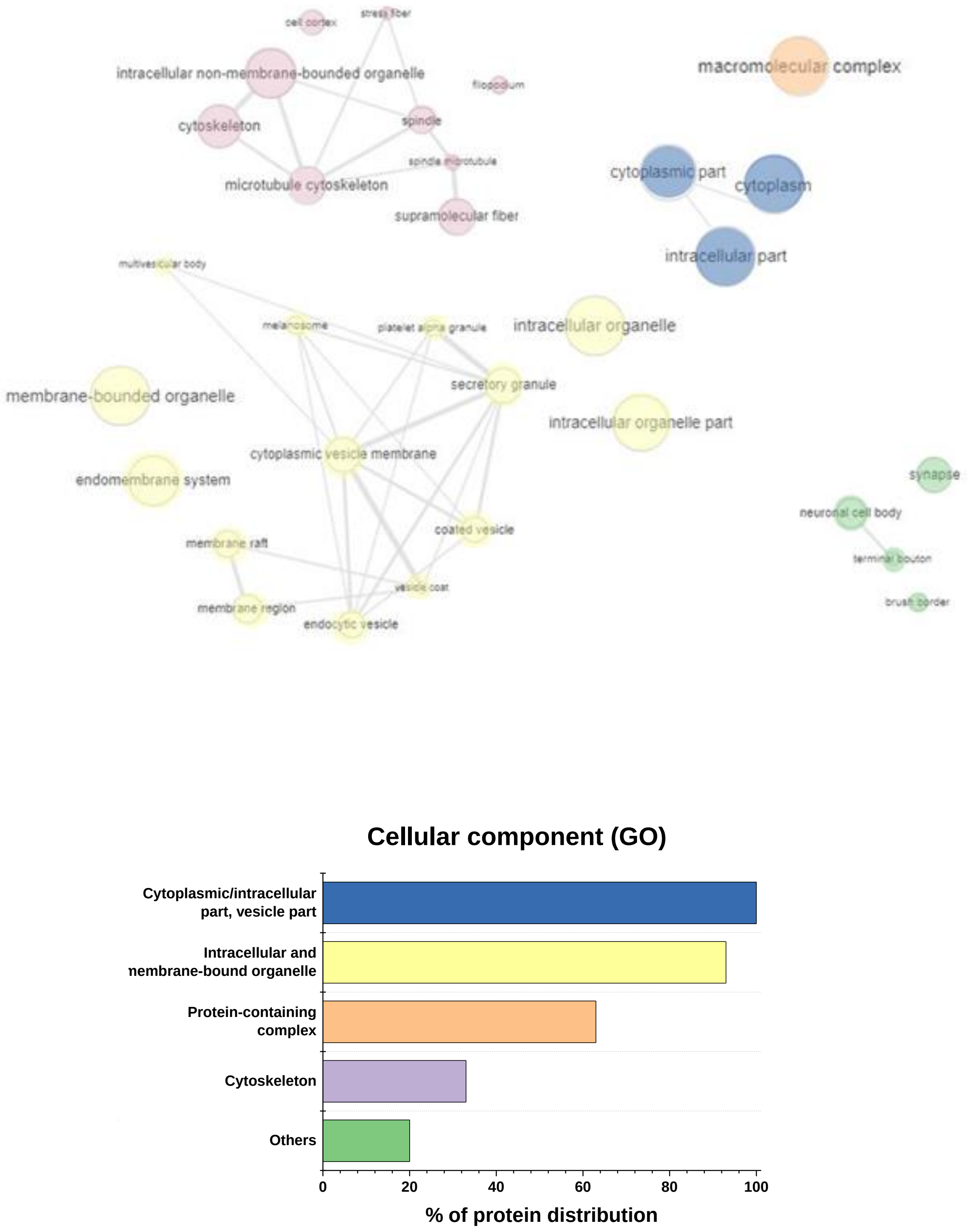


**Fig. S2: GO enrichment analysis elucidating enriched cellular component (CC).** GO terms identified using dataset imported from STRING and analyzed with REVIGO web server. Redundant GO terms were excluded for clarity. Results are visualized with REVIGO interactive maps showing enriched enriched CC GO terms. Similar GO terms are colour coded. The colour matches the bar colours of the bar chart on the right showing percentage.

```
CLUSTAL O(1.2.4) multiple sequence alignment

RFIP3 MASAPPASPPGSEPPGPDPEPGGPDGPGAAQLAPGPAELRLGAPVGGPDPQSPGLDEPAP 60
Rab46 ------------------------------------------------------------

RFIP3 GAAADGGARWSAGPAPGLEGGPRDPGPSAPPPRSGPRGQLASPDAPGPGPRSEAPLPELD 120
Rab46 --------------------------------MAAPDGRV-------------------- 8
                                       :.* *::

RFIP3 PLFSWTEEPEECGPASCPESAPFRLQGSSSSHRARGEVDVFSPFPAPTAGELALEQGPGS 180
Rab46 ------------------VSR-----------------------------PQRLGQGSGQ 21
                         *                                 * ** *.

RFIP3 PPQPSDLSQTHPLPSE--------PVGSQEDGPRLRAVFDALDGDGDGFVRIEDFIQFAT 232
Rab46 GPKGSGA-CLHPLDSLEQKETQEQTSGQLVMLRKAQEFFQTCDAEGKGFIARKDMQRLHK 80
       *: *.    *** *           *.     : : .*:: *.:*.**:  :*: :: .

RFIP3 V--YGAEQVKDLTKYLDPSGLGVISFEDFYQGITAIRN--GDPDGQCYGGVASAQDEEPL 288
Rab46 ELPLSLEELEDVFDALDADGNGYLTPQEFTTGFSHFFFSQNNPSQEDAGEQVAQRHEEKV 140
          . *:::*: . ** .* * :: ::*  *:: :    .:*. :  *  .: :.** :

RFIP3 ACPDEFDDFVTYEANEVTDSAYMGSESTYSECETFTDEDTSTLVHPELQPEGDADSAGGS 348
Rab46 --------YLS---RGDEDLGDMGEDEEA----QF------RMLM---------DRLGAQ 170
              :::   .   * . **.:.       *       ::          *  *..

RFIP3 AVPSECLDAMEEPDHGALLLLPGRPHPHGQSVITVIGGEEHFEDYGEGSEAELSPETLCN 408
Rab46 KVL----E--DESDVKQLWLQLKKEEPHLL---------SNFEDFLTRIISQLQEAHEEK 215
       *     :  :* *   * *   : .**           .:***:     ::*.     :

RFIP3 GQLGCSDPAFLTPSPTKRLSSKKVARYLHQSGALTMEA---LEDPS-------PELMEGP 458
Rab46 NELECALK-------------RKIAAYDEEIQHLYEEMEQQIKSEKEQFLLKDTERFQAR 262
      .:* *:               :*:* * .:   *  *    ::. .        * ::.

RFIP3 EEDIADKVVFLE--------------RRVLELEKDTAATGEQHSRLRQENLQLVHRANAL 504
Rab46 SQELEQKLLCKEQELEQLTQKQKRLEGQCTALHHDKHETKAENTKLKLTNQELA------ 316
      .::: :*::  *               :   *.:*.  *   :::*:  * :*.

RFIP3 EEQLKEQELRACEMVLEET----RRQKELLCKMEREKSIEIENLQTRLQQLDEENSELRS 560
Rab46 -R-----ELERTSWELQDAQQQLESLQQEACKLHQEKEMEVYRVTESL------------ 358
       .     **. .   *:::    .  ::  **:.:**.:*: .:   *

RFIP3 CTPCLKANIERLEEEKQKLLDEIESLTLRLSEEQENKRRMGDRLSHER-HQFQRDKEATQ 619
Rab46 ------------QREKAGLLKQLDFLRERNK-----------HLRDERDICFQKNKAAKA 395
                  :.**  **.::: *  * .           :* .**   **::* *.

RFIP3 ELIE-----------DLRKQLEHLQLLKLEA-----EQRRGRSSSMGLQEYHSRARESEL 663
Rab46 NTAASRASWKKRSGSVIGKYVDSRGILRSQSEEEEEVFGIPRRSSLGLSGYPLTEEEPGT 455
      :               : * ::   :*: ::          * **:**. *    .*

RFIP3 ---------EQEVRRLKQDNR-NL-------------------KEQNEELNGQIITL--- 691
Rab46 GEPGPGGPYPRPLRRIISVEEDPLPQLLDGGFEQPLSKCSEEEEVSDQGVQGQIPEAPPL 515
                : :**: . :.  *                   : .:: ::***

RFIP3 ----------------------SIQGAKSLFSTAFSESLAAEISS----VSRDELMEAIQ 725
Rab46 KLTPTSPRGQPVGKEALCKEESSPSAPDRLFKIVFVGNSAVGKTSFLRRFCEDRFSPGMA 575
                            * .. . **. .* .  *.  :*    ...*.:  .:

RFIP3 KQEEINFRLQD---------------------------YI---DRIIVA-IMETNPSILE 754
Rab46 ATVGIDYRVKTLNVDNSQVALQLWDTAGQERYRCITQQFFRKADGVIVMYDLTDKQSFLS 635
          *::*::                            ::    * :**   :  : *:*.

RFIP3 VK---------------------------------------------------------- 756
Rab46 VRRWLSSVEEAVGDRVPVLLLGNKLDNEKEREVPRGLGEQLATENNLIFYECSAYSGHNT 695
      *:

RFIP3 ------------------------------------
Rab46 KESLLHLARFLKEQEDTVREDTIQVGHPAKKKSCCG 731
```

**Fig. S3: Full Sequence alignment of Rab46 and Rab11FIP3 sequences showing conserved residues involved in dynein binding**. Sequence alignment using Clustal Ω shows a conserved region in the first coiled-coil segment of the dynein adaptor Rab11FIP3 (CC1 box) and Rab46 coil-coiled domain. The alignment also shows a second conserved residue in the spindly motif of RFIP3 (708-712) which is shared within the Rab46 sequence. The two alanines (A435 and A709 in RFIP3) important in the interaction with dynein and the conserved alanines in Rab46 sequence (A227 and A555) are marked in red with red asterisks.

```
CLUSTAL O(1.2.4) multiple sequence alignment

BICD2    ------------------------------------------------------------
spindly  --------------------------------------------------------EADI 4
BIDR2    MSSPDGPSFPSGPLSGGASPSGDEGFFPFVLERRDSFLGGGPGPEEPEDLALQLQ----- 55
HAP1     ------RFVFQGPFGSRATGRG-TGKAAGIWKTPAAYVGRRPGVSGPERAAFIRELEEAL 53
CC_Rab46 ------------------------------------------------------------

BICD2    --------------EVKRLSHELA----------------ETTREKIQAAEYGLAVLEEK 30
spindly  ITN-------------------------------LRCRLKEAEEERLKAAQYGLQLVESQ 33
BIDR2    ----------------------------------------QKEKDLLLAAELGKMLLERN 75
HAP1     CPNLPPPVKKITQEDVKVMLYLLEELLPPVWESVTYGMVLQRERDLNTAARIGQSLVKQN 113
CC_Rab46 ------------------------------------------------IAAYDEEIQHLY 12
                                                          * . : .

BICD2    HQLKLQFEELEVDYEAIRSEMEQLKEAFGQAHTNHKKVAADGESREESLIQESASKEQYY 90
spindly  NELQNQLDKCRNEMMTMTESYEQEKYTLQREVE------------LKSRMLESLSCECEA 81
BIDR2    EELRRQLETLSAQH-LEREE------RLQ--QENH----------ELRRGLAAR--GAEW 114
HAP1     SV-------------------------LM--EENS----------KLEALLGSAKEEILY 136
CC_Rab46 EEMEQQIKSEKEQFLLKDTE------RFQ--ARSQ----------ELEQKLL-------- 46
                                    :

BICD2    VR--------------------KVLELQTELKQLRNVLTNTQSENERLASVAQELKEINQ 130
spindly  IKQQQKMHLEKLEEQLSRSHGQEVNELKTKIEKLKVELDEARLSEKQLK----------H 131
BIDR2    EA--------------------RAVELEGDVEALRAQLGEQRSEQQDSGR---ERARALS 151
HAP1     LR--------------------HQVNLRDELLQLYSDSDEEDEDEEEEEE---E-KEAEE 172
CC_Rab46 ------------------------------------------------------------

BICD2    NVEIQRGRLRDDIKEYKFREARLLQDY--SELEEENISLQKQVSVLRQNQVEFEGLKHEI 188
spindly  QVDH----QKELLSCKSE-ELRVMSERVQESMSSEMLALQIELT-------EMESMKTTL 179
BIDR2    ELSEQNLRLSQQLAQASQ-----TEQ----ELQRELDALRGQCQ-------AQA------ 189
HAP1     EQEE--EEAEEDLQCAHP-----CDAP--KLISQEALLHQHHCP-------QLEALQEKL 216
CC_Rab46 --------------CKEQ-----ELE----QLTQKQKRLEGQCT-------ALH------ 70
                                        : :     . .

BICD2    KRLEEETEYLNSQLEDAIRLKEISERQLE---EALETLKTEREQKNSLRKELSHYMSIND 245
spindly  KEEVNELQYRQEQLELLIT-------NLM---RQVDRLKEEKEERE---KEAVSYYNALE 226
BIDR2    ----L------------------AGAELR---TRLESLQGENQ------------MLQSR 212
HAP1     RLLEE------------------ENHQLREEASQLDTLEDEEQ------------MLILE 246
CC_Rab46 ----H------------------DKHETK---AENTKLKLTNQ----------------- 88
                                   :          *:  .:
```

**Fig. S4: Multiple motif alignment of some dynein adaptors and Rab46 coil-coiled domain**. Sequence alignment using Clustal Ω shows a conserved region (CC1 box) in the first coiled-coil segment of the dynein adaptors BICD2, Spindly, BICDR2, HAP1 and Rab46 coiled-coil domain. Within the conserved region the alanine important in the interaction with dynein and the conserved alanine in Rab46 sequence (A227) are marked in red with red asterisks.

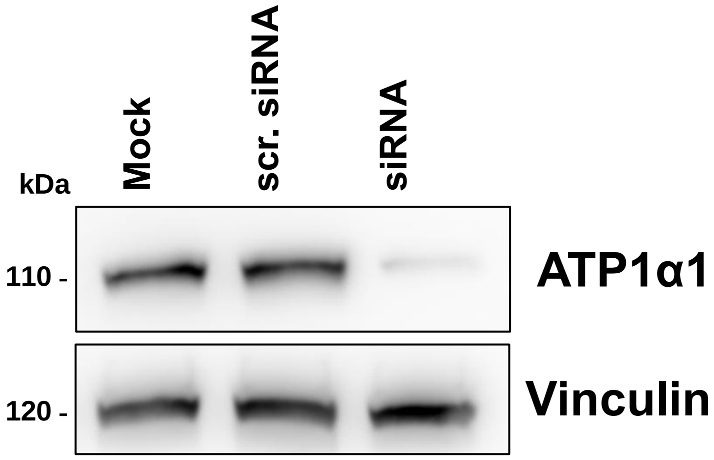


**Fig. S5: Validation of $Na^{2+}$/ $K^{+}$ ATPase subunit α1 antibody.** Representative western blot depicting specificity of ATP1α1 antibody. ATP1α1 depleted endothelial cells, transfected with ATP1α1 siRNA (100 nM), show reduced intensity bands compared to control siRNA. Vinculin used as loading control.
